# Annotation and analysis of *yellow genes* in *Diaphorina citri*, vector for the Huanglongbing disease

**DOI:** 10.1101/2020.12.22.422960

**Authors:** Crissy Massimino, Chad Vosburg, Teresa Shippy, Prashant S. Hosmani, Mirella Flores-Gonzalez, Lukas A. Mueller, Wayne B. Hunter, Joshua B. Benoit, Susan J. Brown, Tom D’Elia, Surya Saha

**Affiliations:** Indian River State College, Fort Pierce, FL 34981; Division of Biology, Kansas State University, Manhattan, KS 66506; Boyce Thompson Institute, Ithaca, NY 14853; USDA-ARS, U.S. Horticultural Research Laboratory, Fort Pierce, FL 34945; Department of Biological Sciences, University of Cincinnati, Cincinnati, OH, 45221; Animal and Comparative Biomedical Sciences, University of Arizona, Tucson, AZ 85721

## Abstract

Huanglongbing (HLB), also known as citrus greening disease, is caused by the bacterium *Candidatus* Liberibacter asiaticus (CLas) and represents a serious threat to global citrus production. This bacteria is transmitted by the Asian citrus psyllid, *Diaphorina citri* (Hemiptera) and there are no effective in-planta treatments for CLas. Therefore, one strategy is to manage the psyllid population. Manual annotation of the *D. citri* genome can identify and characterize gene families that could serve as novel targets for psyllid control. The *yellow* gene family represents an excellent target as *yellow* genes are linked to development and immunity due to their roles in melanization. Combined analysis of the genome with RNA-seq datasets, sequence homology, and phylogenetic trees were used to identify and annotate nine *yellow* genes for the *D. citri* genome. Phylogenetic analysis shows a unique duplication of *yellow-y* in *D. citri*, with life stage specific expression for these two genes. Genomic analysis also indicated the loss of a gene vital to the process of melanization, *yellow-f*, and the gain of a gene which seems to be unique to hemipterans, *yellow 9*. We suggest that *yellow 9* or the gene *yellow 8* (*c*), which consistently groups closely to *yellow-f*, may take on this role. Manual curation of genes in *D. citri* has provided an in-depth analysis of the *yellow* family among hemipteran insects and provides new targets for molecular control of this psyllid pest. Manual annotation was done as part of a collaborative community annotation project (https://citrusgreening.org/annotation/index).

## INTRODUCTION

Huanglongbing, HLB, [Citrus greening disease] is the most serious threat to citrus sustainability [1–4]. HLB is caused by the bacterium *Candidatus* Liberibacter asiaticus, CLas [5]. The Asian citrus psyllid, *Diaphorina citri*, serves as the vector for CLas (Hemiptera:Liviidae) [6]. There is no effective treatment for the pathogen, and pesticide application is the only mechanism to reduce psyllid populations [4]. However current control strategies are not effective and the development of novel mechanisms to reduce the psyllid vector population is necessary. Manual annotation of the *D. citri* genome to identify and characterize genes could identify targets for psyllid control treatments [7–9]. Here, we identified and characterized genes in the *yellow* gene family. The function of *yellow* proteins (dopachrome conversion enzymes, DCE) are involved in the melanin biosynthetic pathway [10]. Melanization is a critical function in insects [11]. Melanization can be triggered locally as an immune effector response by which melanin is synthesized and cross-linked with other molecules in injured areas resulting in the killing of invading pathogens and hardening of the wound clot [12]. Melanization is also essential for cuticle sclerotization or tanning, which leads to hardening of the insect exoskeleton [13], and the prevention of moisture loss [10]. To develop gene-targeting or suppressing treatments for *yellow* proteins, accurate gene sequences needed to be established, annotated, and basic expressional details provided. Here we describe the *yellow* genes of *D. citri*, the Asian citrus psyllid through the combination of genome annotation and expressional differences [14] based on previously conducted RNA-seq studies.

## BACKGROUND

*yellow* genes are of ancient lineage, as evidenced by the presence of *yellow-like* genes in several bacterial species; however, no evidence was found that these genes exist in the complete genome sequences of the worm *Caenorhabditis elegans* or the yeast *Saccharomyces cerevisiae*, suggesting that they may have been lost from many lineages and may now be largely restricted to arthropods [15]. While functional assignments have not yet been made for every member of this family, research suggests that a role in melanization may be conserved for several *yellow* family members [11]. Duplications, as well as losses, are apparent in the *yellow* gene family and phylogenetic analysis shows *yellow* family expansion is associated with insect diversification [11]. Previous studies have shown that the *yellow*-*y, -c, -d, -e, -f, -g*, and -*h* genes were present prior to divergence of the hemimetabolous and holometabolous insects, however, some of these ancestral *yellow* genes are lost in specific insect lineages [11]. The most notable case of *yellow* lineage duplication is the entire Major Royal Jelly Protein (mrjp) family which forms a distinct cluster within the *yellow* family phylogeny and seems to be restricted to certain species of bees (Fig. 1) [15].Here we describe the *yellow* genes of the Asian citrus *psyllid, Diaphorina citri*. Because of the multiplicity of *yellow* genes discovered in *D. citri* and the inconsistency of ortholog names, phylogenetic analysis was performed to properly classify these genes. This was followed by examining expression differences based on previously available RNA-seq datasets. Based on these results, we discuss possible functions of the *yellow* genes identified in D. citri.

**Figure 1:**
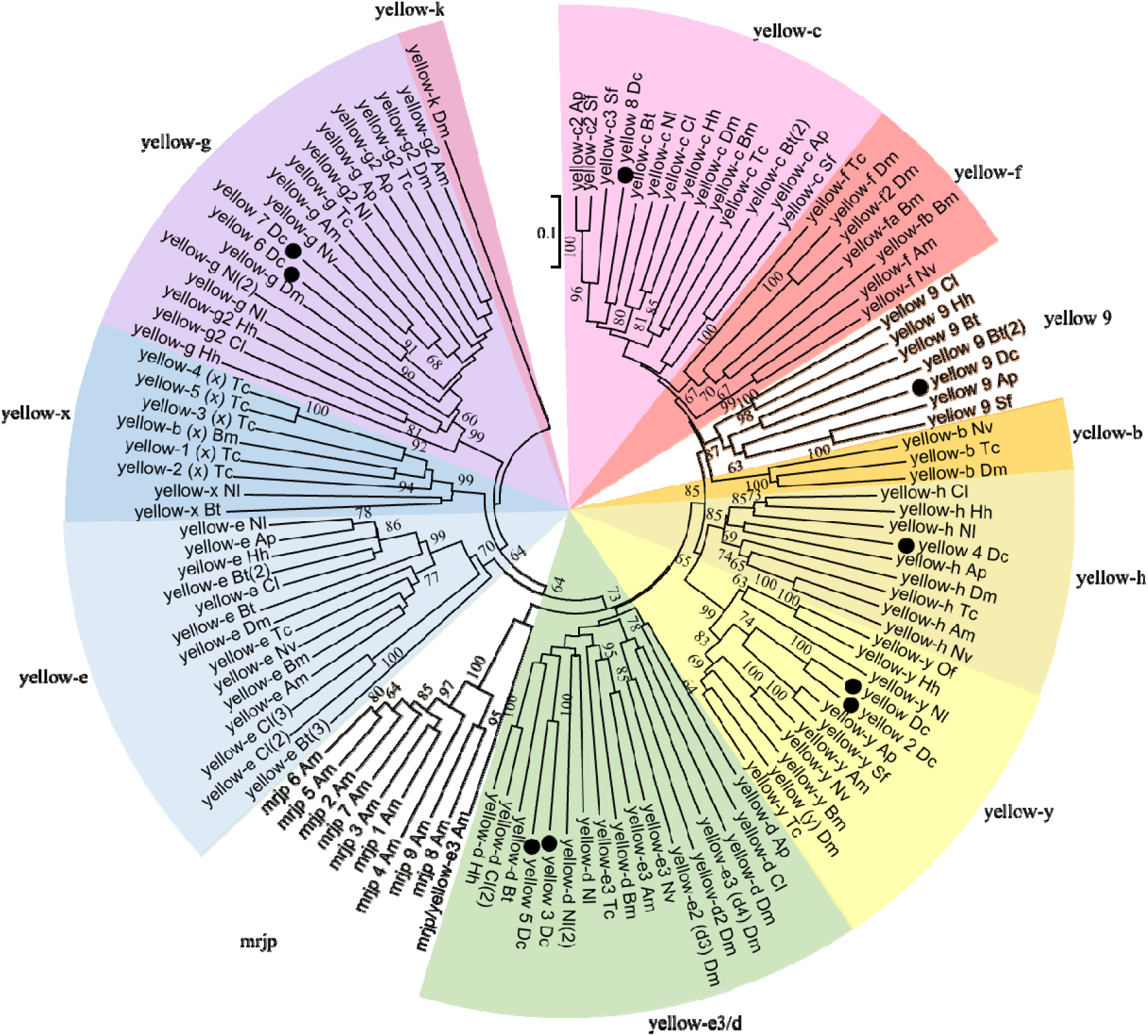
The *yellow* gene family. A neighbor-joining phylogenetic tree was generated from annotated *D. citri* (*Dc*) *yellow* protein sequences and the protein sequence of all predicted *yellow* genes from the insects *Acyrthosiphon pisum* (*Ap*), *Apis mellifera* (*Am*), *Tribolium castaneum* (*Tc*), *Drosophila melanogaster* (*Dm*), *Bombyx mori* (*Bm*), *Sipha flava* (*Sf*), *Bemisia tabaci* (*Bt*), *Nilaparvata lugens* (*Nl*), *Cimex lectularius* (*Cl*), *Halyomorpha halys* (*Hh*). Also included are sequences from Nasonia vitripennis (Nv) and Oncopeltus fasciatus (Of). Bootstrap analysis was performed with 1000 replicates. Values greater than 60 are shown at nodes. *D. citri* sequences are identified with a dot. Color coding indicates specific *yellow* clades. NCBI accession numbers are shown in the Accession Number Table (Table 3, Table 4).

## RESULTS AND DISCUSSION

Manual gene annotation of the *D. citri* genome revealed the presence of nine *yellow* genes, each containing the major royal jelly protein (mrjp) domain conserved in this gene family. All nine of the genes were confirmed by at least four types of evidence that ranged from RNA-seq to ortholog presence (Table 1). Phylogenetic analysis was conducted to determine the orthology of these *yellow* genes and the results coincide well with previous studies (Fig. 1) [11,16]. Based on this analysis, the *yellow* genes in *D. citri* comprise two *yellow*-y genes, two *yellow*-d genes, two *yellow*-*g* genes, one *yellow*-*h*, and one *yellow*-*c*, as well as one *yellow* gene (*yellow* 9) that seems to be a duplication unique to hemipterans, but is closely related to known *yellow*-f orthologs (Fig. 1, Table 2).

**Table 1:** Evidence for gene annotations.

| Gene | Identifier | MCOT | de novo | Iso-Seq | RNA-Seq | Ortholog |
| --- | --- | --- | --- | --- | --- | --- |
| yellow | Dcitr06g10150.1.1 | X | X | X | X |  |
| yellow 2 | Dcitr03g06860.1.1 |  | X | X | X |  |
| yellow 3 | Dcitr07g07190.1.1 | X | X |  | X |  |
| yellow 4 | Dcitr07g07210.1.1 | X | X | X | X | X |
| yellow 5 | Dcitr11g08710.1.1 | X | X |  | X | X |
| yellow 6 | Dcitr02g17890.1.1 | X | X | X | X | X |
| yellow 7 | Dcitr11g06750.1.1 | X | X | X | X | X |
| yellow 8 | Dcitr01g01210.1.1 |  | X | X | X | X |
| yellow 9 | Dcitr01g05760.1.1 | X | X |  | X |  |

**Table 2:** Gene copy numbers.

| Gene | Dc | Ap | Am | Tc | Dm | Bm | Sf | Bt | Nl | Cl | Hh |
| --- | --- | --- | --- | --- | --- | --- | --- | --- | --- | --- | --- |
| yellow-y | 2 | 1 | 1 | 1 | 1 | 1 | 1 | 0 | 1 | 0 | 1 |
| yellow-b | 0 | 0 | 0 | 1 | 1 | 0 | 0 | 0 | 0 | 0 | 0 |
| yellow-f | 0 | 0 | 1 | 1 | 2 | 2 | 0 | 0 | 0 | 0 | 0 |
| yellow-9 | 1 | 1 | 0 | 0 | 0 | 0 | 1 | 2 | 0 | 1 | 1 |
| yellow-c | 1 | 2 | 0 | 1 | 1 | 1 | 3 | 2 | 1 | 1 | 1 |
| yellow-h | 1 | 1 | 1 | 1 | 1 | 0 | 0 | 0 | 1 | 1 | 1 |
| yellow-d/e3 | 2 | 1 | 1 | 1 | 4 | 1 | 0 | 1 | 2 | 2 | 1 |
| yellow-e | 0 | 1 | 1 | 1 | 1 | 1 | 0 | 3 | 1 | 3 | 1 |
| yellow-g | 2 | 2 | 2 | 2 | 2 | 0 | 0 | 0 | 2 | 1 | 2 |
| yellow-x | 0 | 0 | 0 | 5 | 0 | 1 | 0 | 1 | 1 | 0 | 0 |
| yellow-k | 0 | 0 | 0 | 0 | 1 | 0 | 0 | 0 | 0 | 0 | 0 |
| mrjp<sup>†</sup> | 0 | 0 | 10 | 0 | 0 | 0 | 0 | 0 | 0 | 0 | 0 |

Yellow gene family ortholog numbers based on the results from the phylogenetic analysis (Fig.
1). Dc numbers represent the number of manually annotated genes in the *D. citri* v3.0 genome.
<sup>†</sup>Copy number in *Am* includes a predicted sequence from NCBI that groups with mrjp in phylogenetic analysis.

**Table 3:** Non-hemipteran insect accession numbers.

| Gene | *Drosophila melanogaster* | *Tribolium castaneum* | *Bombyx mori* | *Apis mellifera* | *Nasonia vitripennis* |
| --- | --- | --- | --- | --- | --- |
| yellow-y | NP_476792.1 | NP_001161919.1 | NP_001037434.1 | NP_001091693.1 | NP_001154977.1 |
| yellow-b | NP_523586.1 | NP_001161777.1 |  |  | NP_001154989.1 |
| yellow-c | NP_523570.3 | NP_001161778.1 | NP_001037426.1 |  |  |
| yellow-f (fa) † | NP_524335.1 | NP_001161780.1 | NP_001037424.1 | NP_001011635.1 | NP_001154968.1 |
| yellow-f2 (fb)† | NP_650247.1 |  | NP_001037428.1 |  |  |
| yellow-h | NP_651912.3 | XP_008196499.1 |  | NP_001091687.1 | NP_001153394.1 |
| yellow-e | NP_524344.1 | NP_001161779.1 | NP_001159615.1 | XP_003249426.1 | NP_001154985.1 |
| yellow-e2 | NP_650289.2 |  |  |  |  |
| yellow-e3 | NP_650288.1 | NP_001161913.1 |  | NP_001091698.1 | NP_001154982.1 |
| yellow-d | NP_523820.2 |  | NP_001037422.1 |  |  |
| yellow-d2 | NP_611788.1 |  |  |  |  |
| yellow-g | NP_523888.1 | ACY71060.1 |  | XP_396824.1 | XP_008210364.1 |
| yellow-g2 | NP_647710.1 | NP_001161782.1 |  | XP_006559006.1 |  |
| yellow-k | NP_648772.1 |  |  |  |  |
| yellow-1 (x)‡ |  | NP_001161783.1 |  |  |  |
| yellow-2 (x)‡ |  | NP_001161784.1 |  |  |  |
| yellow-3 (x)‡ |  | NP_001161785.1 |  |  |  |
| yellow-4 (x)‡ |  | NP_001161786.1 |  |  |  |
| yellow-5 (x)‡ |  | NP_001165862.1 |  |  |  |
| yellow-b (x)‡ |  |  | NP_001037430.1 |  |  |
| mrjp/yellow-e3 |  |  |  | XP_001122824.1 |  |
| mrjp 1 |  |  |  | NP_001011579.1 |  |
| mrjp 2 |  |  |  | NP_001011580.1 |  |
| mrjp 3 |  |  |  | NP_001011601.1 |  |
| mrjp 4 |  |  |  | NP_001011610.1 |  |
| mrjp 5 |  |  |  | NP_001011599.1 |  |
| mrjp 6 |  |  |  | NP_001011622.1 |  |
| mrjp 7 |  |  |  | NP_001014429.1 |  |
| mrjp 8 |  |  |  | NP_001011564.1 |  |
| mrjp 9 |  |  |  | NP_001019868.1 |  |

Accessions number for non-hemipteran insects used in phylogenetic analysis (Fig. 1). XP
identifiers are computer predicted models. † Yellow-fa and -fb are the names given specifically
to *Bombyx mori*. ‡ Yellow-x is the name commonly used to refer to this group.

**Table 4:** Hemipteran accession numbers.

| Gene | *Acyrtosiphon pisum* | *Nilaparvata lugens* | *Sipha Flava* | *Bemisia tabaci* | *Halymorpha halys* | *Cimex lectularius* | *Oncopeltus fasciatus* |
| --- | --- | --- | --- | --- | --- | --- | --- |
| yellow-y | NP_001165848.1 | XP_022184245.1 | XP_025405013.1 |  | XP_014294277.1 |  | AMW91813.1 |
| yellow-c | XP_008189794.1 | XP_022204541.1 | XP_025415761.1 | XP_018916210.1<br>XP_018902711.1 | XP_014273608.1 | XP_014260013.1 |  |
| yellow-c2 | XP_008189795 |  | XP_025415762.1<br>XP_025421542.1 |  |  |  |  |
| yellow-h | XP_016656930.1 | XP_022203506.1 |  |  | XP_014273783.1 | XP_014247137.1 |  |
| yellow-e | XP_001948479.1 | XP_022204722.1 |  | XP_018907407.1<br>XP_018906454.1<br>XP_018907408.1 | XP_014292044.1 | XP_024083966.1<br>XP_014239663.1<br>XP_014247778.1 |  |
| yellow-d/e3 | XP_001942700.2 | XP_022204983.1<br>XP_022205442.1 |  | XP_018916419.1 | XP_014275903.1 | XP_024083964.1<br>XP_024083953.1 |  |
| yellow-g | XP_001944949.2 | XP_022207733.1<br>XP_022191720.1 |  |  | XP_014292984.1 |  |  |
| yellow-g2 | XP_001945004.2 | XP_022205647.1 |  |  | XP_014292982.2 | XP_014244848.1 |  |
| yellow-x |  | XP_022206946.1 |  | XP_018912253.1 |  |  |  |
| yellow 9 | XP_001946728.1 |  | XP_025413197.1 | XP_018905078.1<br>XP_018901286.1 | XP_014288983.1 | XP_014255102.1 |  |

Accession numbers for hemipterans used in phylogenetic analysis (Fig. 1). XP identifiers are computer predicted models.

There are nine *yellow* genes in *D. citri* based on transcript and ortholog evidence (Table 1). Each gene has been assigned an OGS3 identifier. Evidence from a *de novo* Oases or Trinity model from an independent transcriptome known as MCOT [9] was used to validate or improve our models. Illumina (RNA-Seq) reads were used to help create or validate our model. A de novo transcriptome was built from long RNA-Seq reads generated with Pacific Biosciences technology (Iso-Seq) and was used to help validate the exon structure of the models. These full length transcripts allowed us to disambiguate noisy signals from short read RNAseq data. Orthologous sequences from related insects and information about conserved motifs or domains were used to determine the final annotation. We used proteins from *Drosophila melanogaster* (Dm) [17], *Tribolium castaneum* (*Tc*) [18,19], *Bombyx mori* (*Bm*) [20], *Apis mellifera* (*Am*) [21], *Nasonia vitripennis* (*Nv*) [22], *Acyrthosiphon pisum* (*Ap*) [23], *Nilaparvata lugens* (*Nl*) [24,25], *Sipha flava* (*Sf*) [26], *Bemisia tabaci* (*Bt*) [27], *Halyomorpha halys* (*Hh*) [28], *Cimex lectularius* (Cl) [29], and *Oncopeltus fasciatus* (Of) [30].

Table 2 lists yellow gene family ortholog numbers in *A. pisum, A. mellifera, T. castaneum, D. melanogaster, B. mori, S. flava, B. tabaci, N. lugens, C. lectularius, and H. halys*. Genes manually annotated in the *D. citri* v3.0 genome are listed in the column labeled Dc. Phylogenetic analysis was performed using sequences in tables 3 and 4 (Fig. 1). *yellow*-f was not found in the genome or *de novo* transcriptome of *D. citri*, or in the hemipteran sequences used in this analysis. Instead, hemipterans formed a separate clade which is labeled as *yellow* 9 here (Table 2, Fig. 1).

### yellow-y

*yellow*-y (also simply referred to as *yellow*) was the first example of a single gene mutation affecting behavior [13]. *yellow*, however, was initially identified because of its role in pigmentation and was named for the loss of black pigment that gave mutant flies a more *yellow* appearance [31]. Recent studies suggest that the *Drosophila melanogaster yellow*-*y* and ebony gene together determine the pattern and intensity of melanization [32], and that the *yellow*-y gene may regulate the expression of *yellow*-f or other enzymes involved in melanization [10]. Many studies have also noted a role for *yellow*-y in the behavior and mating ability of Drosophila, such as changes in the structures used during mating in *yellow* mutants [13]. *yellow*-y is present in most insect species as a single copy, however, both *yellow* and *yellow* 2 in *D. citri* form a clade exclusively with known *yellow*-y orthologs, indicating a duplication event that seems to be unique to *D. citri* (Fig. 1). These two *yellow*-*y* genes show inverse expression patterns in D. citri; that is, *yellow* (y) shows highest expression in the nymph, while *yellow 2* (*y2*) shows highest expression in the adult (Fig. 2). The high expression of *yellow* (y) in nymphs correlates with research that has found *yellow*-y to be abundant in *Drosophila* pupae, when melanin is deposited in the adult cuticle [33]. *yellow* 2 (*y2*), on the other hand, may play an important role in adult *D. citri* and should be studied further.

**Figure 2:**
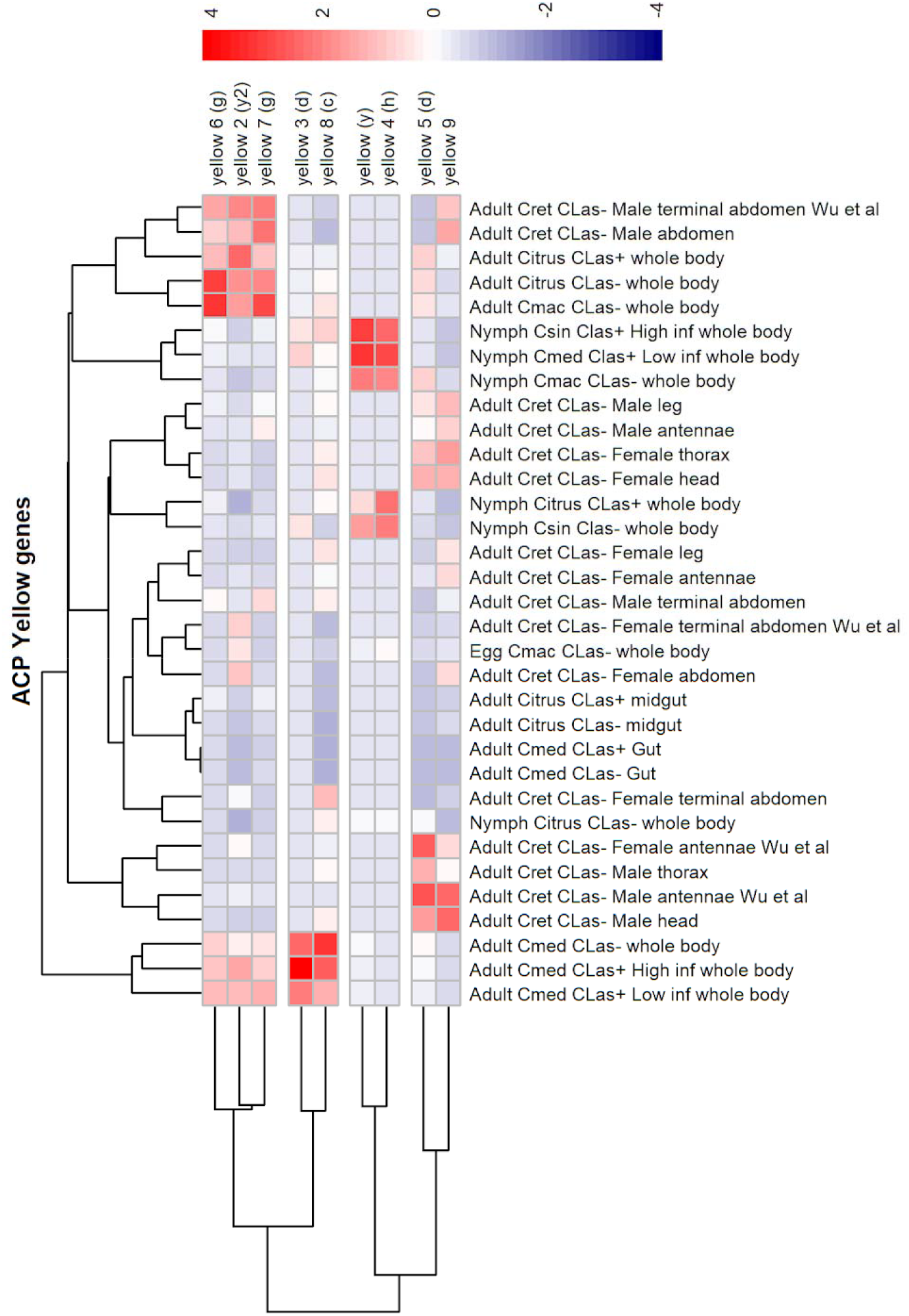
Comparative expression levels of the *D. citri yellow*-y proteins in infected vs. uninfected *D. citri* insects grown on various citrus varieties. Expression data were obtained using the Citrus Greening Expression Network (http://cgen.citrusgreening.org/) [14]. RNA-Seq data for psyllids were obtained from NCBI BioProject’s PRJNA609978 and PRJNA448935 in addition to published data sets [8,34–37]. Citrus hosts are abbreviated as Cmed (C. medica), Cret. (*C. reticulata*) and Cmac (*C. macrophylla*).

### yellow-c, -f, -b, and 9

*yellow*-*b* and *yellow*-*c* are *yellow* genes to which no function has been directly identified. Phylogenetic analysis does, however, reveal a close relationship between *yellow*-*b, yellow*-*c, yellow*-*f*, and *yellow* 9 [11,16] (Fig. 1). *yellow*-*f* and *yellow*-*f2* in Drosophila have been found to function as dopachrome conversion enzymes (DCE) that likely play an important role during melanin biosynthesis [32]. Interestingly, while most hemipterans seem to have one or more *yellow*-*c* genes, none form clades exclusively with known *yellow*-*f* or *yellow*-*b* orthologs (Fig. 1). Instead, hemipteran genes are grouped into their own separate clade. A close relationship was observed, however, between *yellow*-*f* and *yellow* 9, which is supported by the presence of an Acyrthosiphon pisum sequence in the *yellow* 9 clade, previously reported as grouping with *yellow*-*f* (Fig. 1) [11]. The addition of several other hemipteran sequences may have helped align *A. pisum* more closely to the *yellow* 9 orthologs. This distinctness of the hemipteran group is common among the other *yellow* genes in the tree, however, the association to a known ortholog is typically much clearer than is seen with *yellow* 9. More studies should be conducted to conclusively determine the true identity of this hemipteran outlier.

*yellow* 8 (*c*) shows the greatest expression levels of all *yellow* genes in *D. citri* and is most highly expressed in the adult whole body of *D. citri* insects reared on Citrus medica (Fig. 2). There was a significant increase in expression in nymphs reared on CLas positive Citrus sinensis, with 3.85-fold upregulation in whole nymphs (Fig. 3a). The gene was also upregulated by 3.75-fold in the midgut of infected adult psyllids reared on Citrus spp. (Fig. 3b). These results suggest there may be an interaction between *yellow* 8 (*c*) and pathogen infection, therefore warranting further investigations.

**Figure 3:**
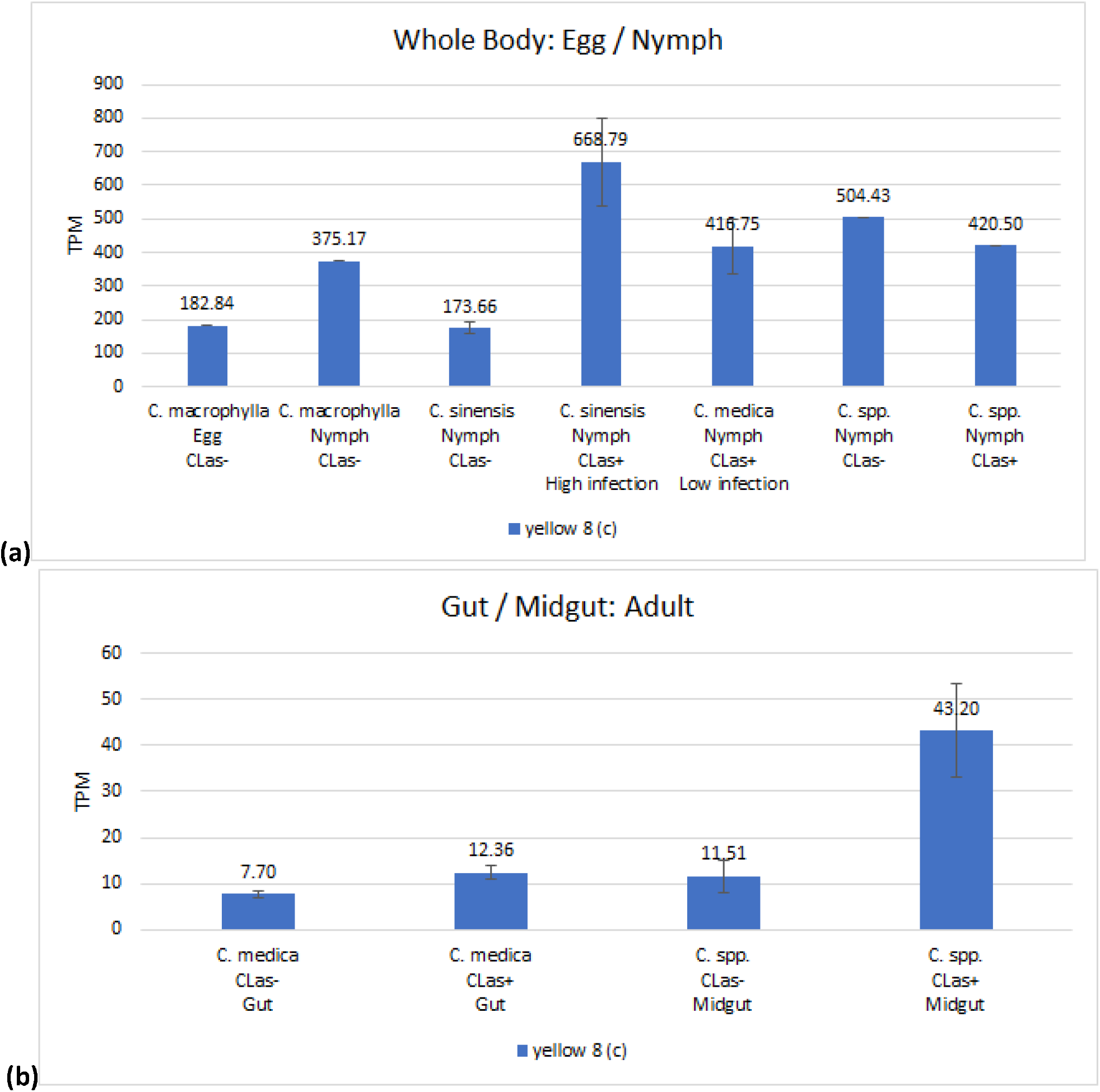
Comparative expression levels of the *D. citri* yellow 8 (c) proteins in uninfected *D. citri* insects grown on various citrus varieties. Values are represented in transcripts per million (TPM). Expression data obtained using the Citrus Greening Expression Network (http://cgen.citrusgreening.org/) [14]. **(a)** Comparative expression levels in the whole body of *D. citri* egg and nymphs. Eggs and nymphs raised on Citrus macrophylla [8]and nymphs raised on Citrus spp. [35] are single replicate data. RNA-Seq data were sourced from insects raised on *C. sinensis* and *C. medica* (NCBI BioProject PRJNA609978). **(b)** Comparative expression levels in the gut [34] and midgut [36] of *D. citri* adults.

### yellow-h

*yellow*-*h* transcripts show color-related expression patterns in some species, but the function of the encoded protein is poorly understood [11]. Phylogenetic analysis reveals that *D. citri* contains one *yellow*-*h* gene, *yellow* 4 (Table 2, Fig. 1). Expression data from *D. citri* shows the highest expression of this gene in the egg and nymph (Fig. 2). This is consistent with previous research showing mutations of *yellow*-*h* in larval Vanessa cardui led to death in pupal stages of development, suggesting that *yellow*-*h* could be important during insect development [16]. Furthermore, expression data revealed an 8.47-fold increase in *yellow*-*h* expression in *D. citri* nymphs reared on *Citrus spp*. and infected with CLas (52.52 TPM) versus uninfected nymphs (6.2 TPM) (Fig. 4). This differential expression of *yellow*-*h*, coupled with the impact of mutations in pupal mortality, indicates that *yellow*-*h* could be a potential RNAi target and warrants additional study in *D. citri*.

**Figure 4:**
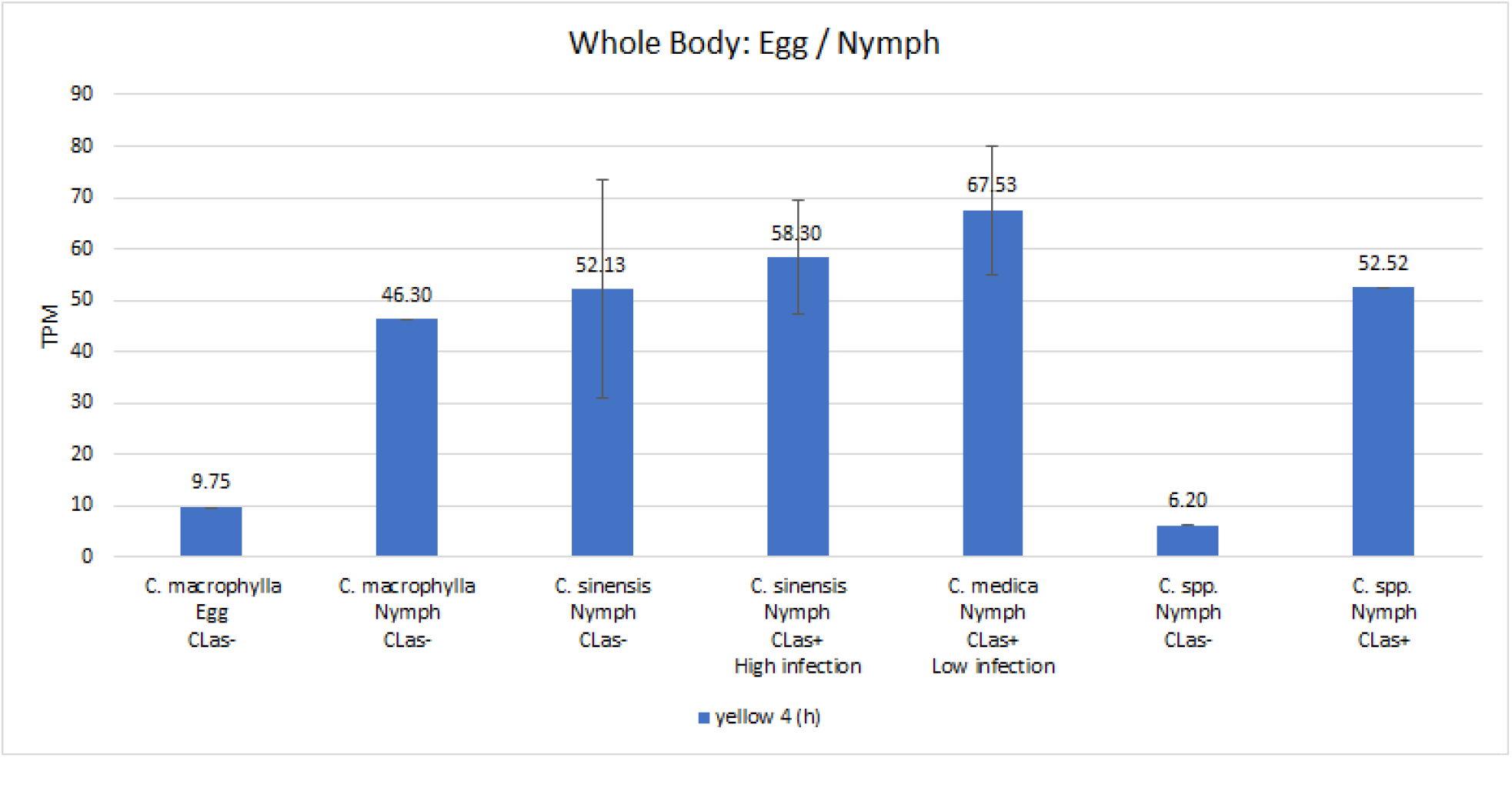
Comparative expression levels of the *D. citri yellow* 4 (h) proteins in eggs, nymphs and whole body infected vs. uninfected *D. citri* adults grown on various citrus varieties. Values are represented in transcripts per million (TPM). Eggs and nymphs raised on Citrus macrophylla [8]and nymphs raised on Citrus spp. [35] are single replicate data. RNA-Seq data were sourced from insects raised on C. sinensis and *C. medica* (NCBI BioProject PRJNA609978). Expression analysis was performed using the Citrus Greening Expression Network (http://cgen.citrusgreening.org/) [14].

### yellow-e3/d

Previous research has revealed that *yellow*-d shows red-specific expression in the butterfly *V. cardui*, and that the loss of *yellow*-d function not only affects melanin patterns, but also presumptive ommochrome patterns [16]. Phylogenetic analysis of the *yellow* genes annotated in the *D. citri* genome shows *yellow* 3 and *yellow* 5 in a clade with known *yellow*-e3/d orthologs (Fig. 1). Expression of *yellow* 3(d) was highest in the whole body of adult psyllid reared on *Citrus medica*, while those reared on *C. macrophylla* or C. spp. showed low expression; and similarly, expression was highest in nymphs raised on *C. sinensis* and *C. medica*, with almost no expression in psyllids reared on other citrus species (Fig. 2). In insects reared on *C. reticulata*, expression of *yellow*3(*d*) was consistently very close to zero TPM while *yellow* 5(*d*), on the other hand, showed relatively high expression in the adult antennae, head, and thorax (Fig. 5).

**Figure 5:**
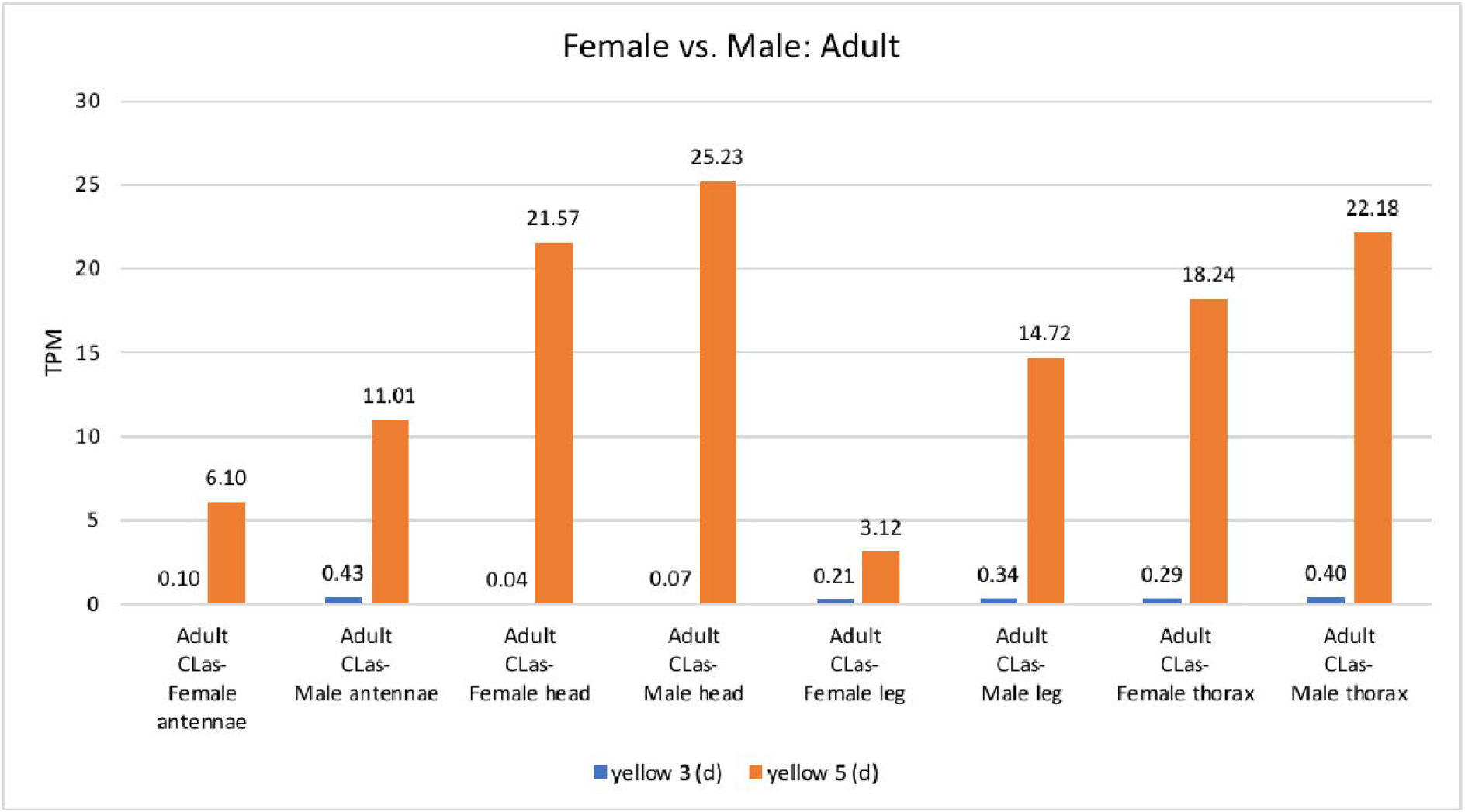
Comparative expression levels of the *D. citri yellow*-d proteins in infected vs. uninfected *D. citri* insects grown on *C. reticulata*. Expression data were obtained using the Citrus Greening Expression Network (http://cgen.citrusgreening.org) [14]. These samples from psyllid tissues have a single replicate and are from NCBI BioProject PRJNA448935.

### yellow-g

The function of *yellow*-g is currently not well understood, however, it is often present in duplicate in most insects (Fig. 1, Table 2). *D. citri* contains two *yellow*-g genes, *yellow* 6 and *yellow* 7, both of which are expressed more in adult males versus females raised on C. reticulata (Fig. 2). The expression of both genes is relatively similar throughout the stages and tissues that have been assayed. Neither gene is expressed in the egg, and both show low expression levels in the nymph with higher expression in adults. There is a notable upregulation of *yellow 6* (*g*) from undetectable in uninfected nymphs raised on C. spp. to 4.01 TPM in infected nymphs. There is also an upregulation of *yellow 6* (*g*) by 8-fold and *yellow* 7 (g) by 2.64- fold in the midgut of infected versus uninfected adult psyllids reared on *C. spp*. (Fig. 6). This effect may indicate an immune response and should be studied further as a possible RNAi target.

**Figure 6:**
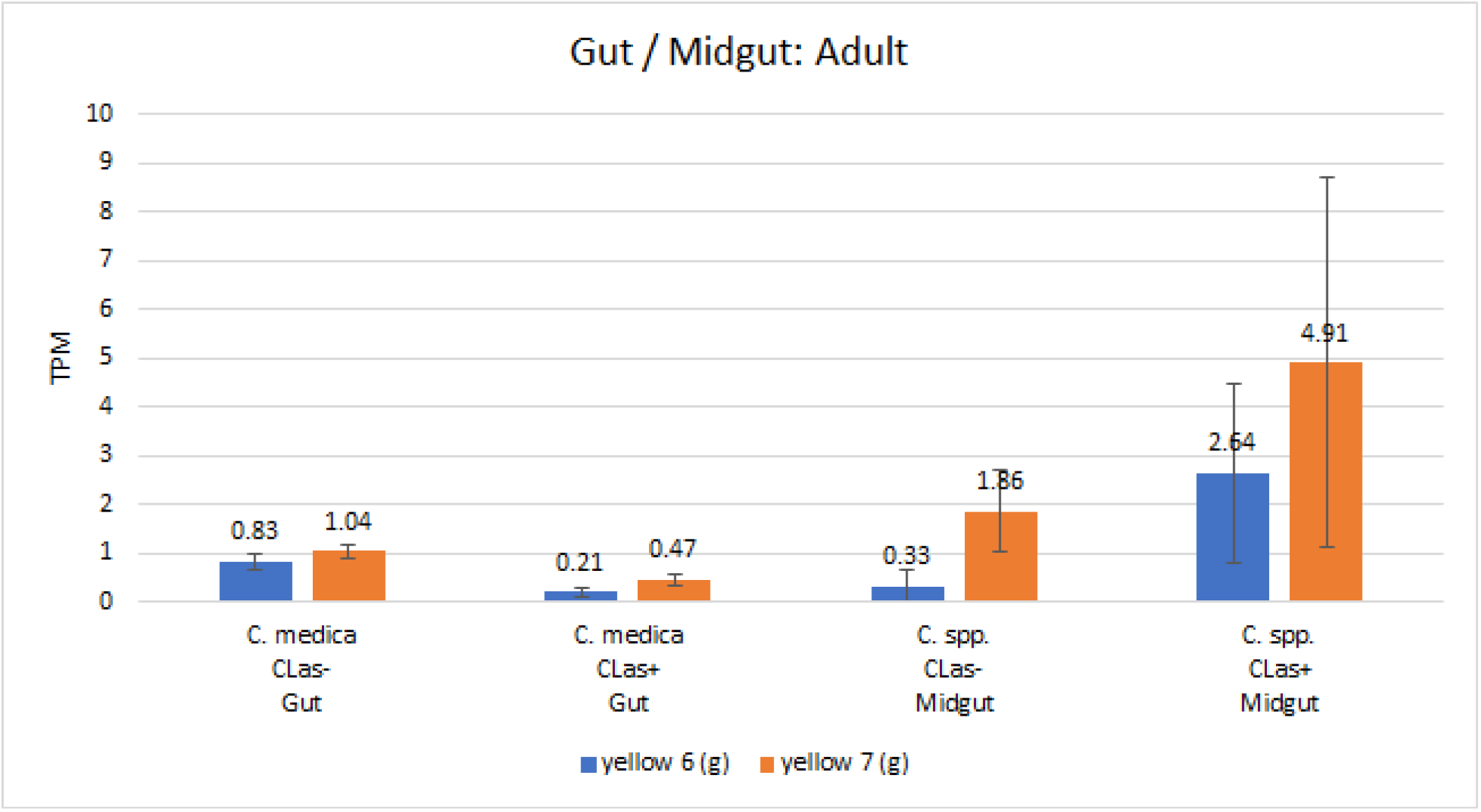
Comparative expression levels of the *D. citri* yellow 6 and yellow 7 (d) proteins in *D. citri* gut [34] and midgut [36] of infected and uninfected adult insects. Values are represented in transcripts per million (TPM). Expression data were obtained using the Citrus Greening Expression Network (http://cgen.citrusgreening.org) [14].

## CONCLUSION

The yellow gene family is a continuously evolving set of genes, with duplications and losses among insects [11]. Many of these genes are crucial in the process of melanization, which is essential for insect survival in relation to development and immunity [10,12]. Though the function of some *yellow* proteins are still not well understood, identification of these in the hemipteran, *D. citri*, provides a novel insect lineage for studies on insect evolution and biology. *D. citri* harbors a unique duplication of *yellow*-y, a gene which may affect cuticular hardening and, therefore, could be a potential target for a *D. citri*-specific molecular control mechanism [13]. Expression data shows an inverse relationship between the two *yellow*-y genes, suggesting independent roles for these proteins during juvenile and adult stages (Fig. 2). The *yellow* 9 gene appears to be unique to hemipterans (Fig. 1), and is a potential alternative to *yellow*-f in holometabolus insects which encodes for dopachrome conversion enzyme (DCE) in Drosophila [32]. Future directed studies are required to confirm the role of *yellow*-9. Continued examination of the *yellow* gene family across arthropods, and especially in insect vectors like *D. citri*, provide novel and species-specific gene targets, potentially through the use of RNA interference, to control psyllid populations and reduce the impacts of pathogens, such CLas causing citrus greening.

## METHODS

The *D. citri* genome was annotated as part of an annotation community driven strategy [9] through an undergraduate-focused program [38]. Protein sequences of the *yellow* family were collected from the NCBI protein database and were used for a blast search of the *D. citri* MCOT protein database [39]. The MCOT protein sequences were used to search the *D. citri* genomes (version 2.0 and 3.0). The regions of high sequence identity were manually annotated in Apollo, using de novo transcriptome and MCOT gene predictions, RNA-Seq, Iso-Seq, and ortholog data as evidence to determine and validate proper gene structure (Table 1). The gene models were compared to those from other hemipterans for accuracy and completeness. A neighbor-joining phylogenetic tree of *D. citri yellow* protein sequences along with was created in MEGA version 7 using the MUSCLE multiple sequence alignment with p-distance for determining branch length and 1,000 bootstrap replicates [40]. A more detailed description of the annotation workflow is available via protocols.io [41]. Accession numbers for the sequences used in this analysis can be found in tables 1,3, and 4 and in the supplementary files. Comparative expression levels of *yellow* proteins throughout different life stages (egg, nymph, and adult) in Candidatus Liberibacter asiaticus (*C*las) exposed vs. healthy *D. citri* insects was determined using RNA-seq data and the Citrus Greening Expression Network (http://cgen.citrusgreening.org). Gene expression levels were obtained from the Citrusgreening Expression Network [14] and visualized using Excel and the pheatmap package in R [42,43].

## Abbreviations

*Am*: *Apis mellifera*
*A. mellifera*: *Apis mellifera*
*A. pisum*: *Acyrthosiphon pisum*
*Ap*: *Acyrthosiphon pisum*
BLASTp: *protein BLAST*
*Bm*: *Bombyx mori*
*Bt*: *Bemisia tabaci*
*Cl*: *Cimex lectularius*
*C*Las: *Candidatus* Liberibacter asiaticus
DCE: dopachrome conversion enzyme
*Dc*: *Diaphorina citri;*
*Dm*: *Drosophila melanogaster*
*Hh*: *Halyomorpha halys*
Iso- Seq: Isoform sequencing
MCOT: Maker, Cufflinks, Oasis, Trinity
mrjp: major royal jelly protein
NCBI: National Center for Biotechnology Information
*Nl*: *Nilaparvata lugens*
*N. vitripennis*: *Nasonia vitripennis*
PCR: polymerase chain reaction
CGEN: Citrus Greening Expression Network
RNAi: RNA interference
RNA-Seq: RNA sequencing
*Sf*: *Sipha flava*
sgRNA: single guide RNA
*Tc*: *Tribolium castaneum*
TPM: Transcripts per million
*V. cardui*: *Vanessa cardui*

## Authors’ contributions

Crissy Massimino performed investigation of research, visualization of data, and writing of original draft. Chad Vosburg provided supervision and validation of research.

## Acknowledgements

We would like to thank Helen Wiersma-Koch (Indian River State College), Thomson Paris (USDA-ARS-Horticultural Research Laboratories) for assistance.

